# The trypanocidal benzoxaborole AN7973 inhibits trypanosome mRNA processing

**DOI:** 10.1101/295550

**Authors:** Daniela Begolo, Isabel M Vincent, Federica Giordani, Michael J Witty, Timothy G Rowan, Zakaria Bengaly, Kirsten Gillingwater, Yvonne Freund, Michael P Barrett, Christine Clayton

## Abstract

Kinetoplastid parasites - trypanosomes and leishmanias - infect millions of humans and cause economically devastating diseases of livestock, and the few existing drugs have serious deficiencies. Benzoxaborole-based compounds are very promising potential novel anti-trypanosomal therapies, with candidates already in human and animal clinical trials. Their targets in trypanosomes were hitherto unknown. We investigated the mechanism of action of several benzoxaboroles, including AN7973, an early candidate for veterinary trypanosomosis.

In all kinetoplastids, transcription is polycistronic. Individual mRNA 5’-ends are created by *trans* splicing of a short leader sequence, with coupled polyadenylation of the preceding mRNA. Treatment of *Trypanosoma brucei* with AN7973 inhibited *trans* splicing within 1h, as judged by loss of the Y-structure splicing intermediate and reduced levels of mRNA, and accumulation of peri-nuclear granules which are typical for splicing inhibition. Methylation of the spliced leader precursor RNA was not affected, but more prolonged AN7973 treatment caused an increase in S-adenosyl methionine and methylated lysine. Together, these results indicate that mRNA processing is the primary target of AN7973. Polyadenylation is required for kinetoplastid *trans* splicing. The EC_50_ for AN7973 in *T. brucei* was increased three-fold by over-expression of the *T. brucei* cleavage and polyadenylation factor CPSF3, identifying CPSF3 as a potential molecular target. Our results thus chemically validate mRNA processing as a viable drug target in trypanosomes.

Several other benzoxaboroles showed metabolomic and splicing effects that were similar to those of AN7973, identifying splicing inhibition as a common mode of action, and suggesting that it might be linked to subsequent changes in methylated metabolites. Granule formation, splicing inhibition, and resistance after CPSF3 expression did not, however, always correlate, and prolonged selection of trypanosomes in AN7973 resulted in only 1.5-fold resistance. This suggests that the modes of action of oxaboroles that target trypanosome mRNA processing may extend beyond CPSF3 inhibition.

**Author summary:** Trypanosomes and leishmanias infect millions of humans and cause economically devastating diseases of livestock; the few existing drugs have serious deficiencies. Trypanosomosis of cattle, caused mainly by *Trypanosoma congolense* and *Trypanosoma vivax*, is a serious problem in Africa, because bovids are used not only for meat and milk, but also for traction. Only two drugs are in routine use for chemotherapy and chemoprophylaxis of bovine trypanosomosis. A single injection of the benzoxaborole compound AN7973 was sufficient to cure *T. congolense* infection in cattle and goats, but AN7973 was less effective against *T. vivax*. This precluded development of AN7973 as a commercially viable treatment against cattle trypanosomosis, but it could still have potential for diseases caused by other salivarian trypanosomes.

We used a large range of methods to find out how AN7973 kills trypanosomes, and compared it with several other benzoxaboroles. AN7973 and some of the other compounds had effects on parasite metabolism that resembled those previously seen for a benzoxaborole that is being tested for human sleeping sickness. The most rapid effect of AN7973, however, was on processing of trypanosome mRNA. As a consequence, amounts of mRNA decreased and synthesis of proteins stopped. We conclude that AN7973 and some other benzoxaboroles kill trypanosomes by stopping gene expression.

## Introduction

Kinetoplastid protists cause severe human diseases affecting millions of people. *Trypanosoma cruzi* causes Chagas disease in South America, and various *Leishmania* species cause a spectrum of diseases throughout the tropics. Salivarian trypanosomes, the subject of this study, cause sleeping sickness in humans and economically important diseases in cattle, horses and camels [1-3]. Approximately 70 million people, living in sub-Saharan Africa, are estimated to be at risk of contracting human African trypanosomosis, which is caused by *Trypanosoma brucei* subspecies [4, 5]. As a result of sustained international activities to control the disease [4-6], less than 3000 cases were reported in 2016 (http://www.who.int/trypanosomiasis_african/en/). Trypanosomosis in cattle, caused by infection with *Trypanosoma congolense*, *Trypanosoma vivax* and, to a lesser extent, *T. brucei*, is in contrast a major problem, with wide-reaching effects on human well-being: cattle are used not only as a source of milk and meat, but also for traction. Elimination of cattle trypanosomosis could create economic benefits estimated at nearly 2.5 billion US$ per year [2]. Within Africa, trypanosomosis is transmitted by tsetse flies, but outside Africa, variants of *T. brucei* are transmitted venereally or by biting flies, and *T. vivax* can also be transmitted non cyclically by non-tsetse biting flies with massive economic losses affecting draught and milk animals from Argentina to the Philippines [7].

Control of cattle trypanosomosis currently relies on reducing the tsetse population by means of traps and insecticidal dips, together with treatment as required. The most popular treatment is with the diamidine diminazene aceturate (Berenil), the alternative being the DNA-intercalating molecule isometamidium [3]. Suramin is also sometimes used [3]. Development of new animal therapeutics is constrained by the need for cure after a single intramuscular injection [3].

Early human trypanosomosis prior to CNS involvement is treated with pentamidine (another diamidine) or suramin, while the late-stage disease, with CNS involvement, has to be treated with either arsenicals or a combination of eflornithine (difluoromethyl ornithine, DFMO) and nifurtimox [8, 9]. These drugs are up to a century old and suffer from numerous disadvantages including severe toxicity and emerging resistance [8, 9]. Recent efforts to develop new therapies have been supported mainly by product development partnerships with both public and private funding, and these are beginning to show some success. Fexinidazole, for example, is a nitrohetrocyclic compound that has recently completed phase 2/3 trials [10]. Although fexinidazole requires a 10 day treatment and must be administered with food, as a once a day oral treatment it has the potential to dramatically simplify treatment. The current standard of care for stage 2 HAT is NECT (nifurtimox-eflornithine combination therapy) which requires 14 slow intravenous infusions of eflornithine of 2 hours each over 7 days, together with three times a day oral nifurtimox for 10 days. Treatment with NECT requires specialized hospital administration and trained staff.

In the last ten years, benzoxaboroles have generated considerable excitement for antimicrobial and other applications. Benzoxaboroles have a range of known effects. For example, they are able to bind *cis*-diols, such as those found in sugars, yielding stable spiro complexes [11]. This activity is the basis of the mode of action of the antifungal drug Tavaborole (AN2690) [12]. AN2690 binds to the editing site of leucyl tRNA synthetase, forming an adduct with the 3’-terminal adenosine of tRNA^Leu^ via two of the ribose hydroxyl groups [13]. This traps the tRNA in the editing site, preventing further function of the enzyme. Binding of AN2690 is stabilised, and therefore made specific, by interactions of other parts of the molecule with the enzyme polypeptide. Other oxaboroles inhibit gram-negative and mycobacterial leucyl tRNA synthetases by the same mechanism [14-16]. The binding of oxaboroles is, however, not confined to the cis-diol mode, or to interaction with sugars. Two classes, for example, interact with ATP-binding pockets, but with different modes of binding. Crisaborole is approved for the treatment of atopic dermatitis [17, 18]. It selectively inhibits phosphodiesterase PDE4, with the boron being coordinated with zinc and magnesium ions within the active site [19]. In contrast, for aminomethylphenoxy benzoxaboroles that inhibit Rho-activated protein kinases the pharmacophore provides hydrogen bond donors and acceptors to the hinge and the aminomethyl group interacts with the magnesium/ATP-interacting aspartic acid. [20].

The first trypanocidal oxaboroles to be described were the oxaborole 6-carboxamides [21, 22]. Acoziborole (=SCYX-7158 or AN5568, Fig 1) has good pharmacokinetics and efficacy in mouse trypanosomosis [23]. Its features include oral availability and the ability to cross the blood brain barrier, both of which are very important for drugs against the human disease [21], and it is now in phase IIb/III clinical trials for sleeping sickness (see https://www.dndi.org/diseases-projects/portfolio/). AN7973 (Fig 1), the main subject of this paper, is efficacious against *T. congolense* and *T. brucei*, and was considered as a candidate for treatment of cattle trypanosomosis, but was later replaced by AN11736 (Fig 1), which can achieve single-dose cure of both *T. congolense* and *T. vivax* infection in cattle [24].

**Fig 1.**
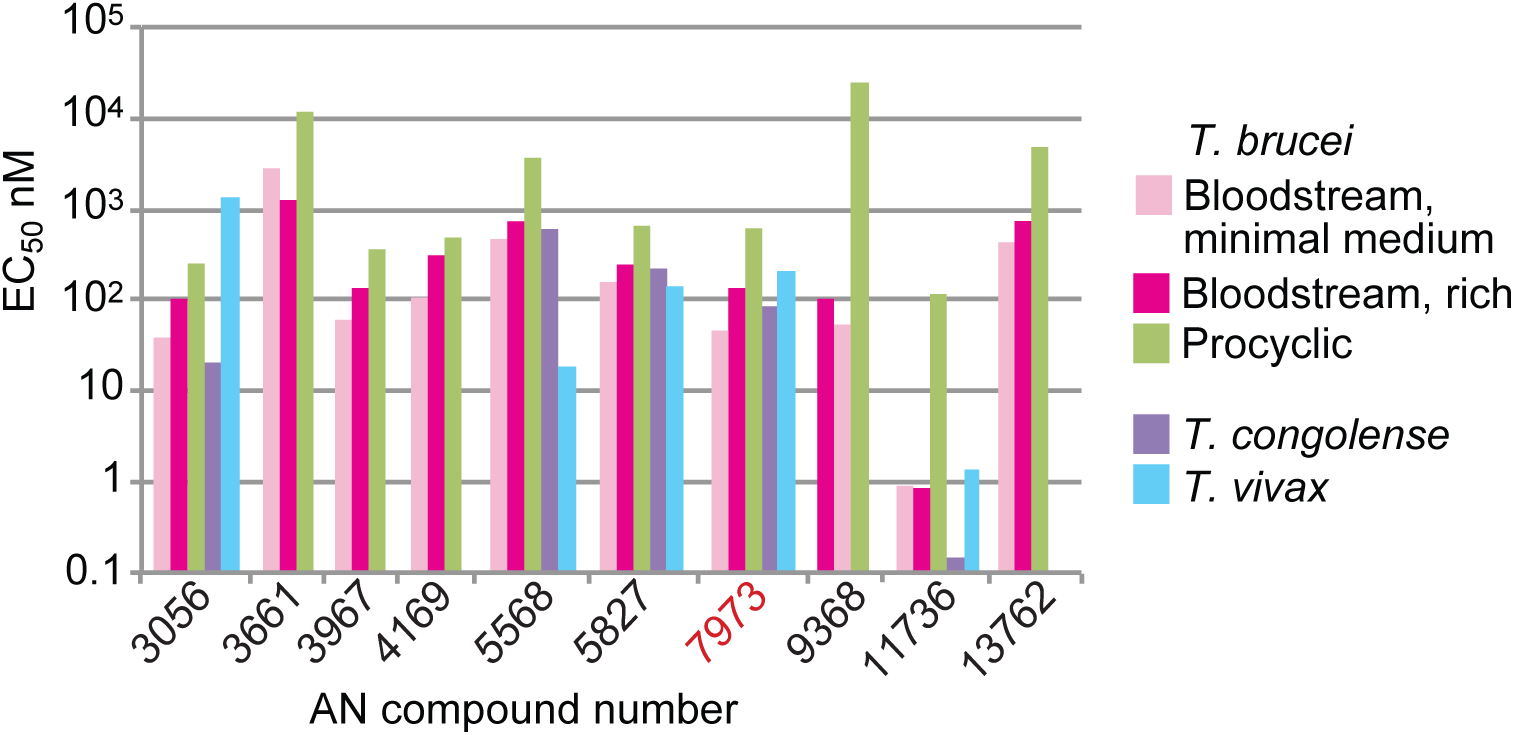
EC_50_s of selected benzoxaboroles for salivarian trypanosomes. The graph shows EC_50_s against *T. brucei* (in vitro, 48h assay for bloodstream forms, 72h for procyclic forms) and for selected compounds, also against *T. congolense* (*in vitro*) and *T. vivax* (*ex vivo*). Details for all tested compounds are in S1 Table.

At present, not enough is known about structure-activity relationships in benzoxaboroles to enable general predictions to be made about their modes of action, and experimental observations have not revealed direct targets in trypanosomes. Jones et al. [25] obtained three independent *T. brucei* lines with 4-5-fold resistance upon 6-month selection with increasing levels of AN2965 (Fig 1, named oxaborole-1 in their paper). They observed numerous genome rearrangements and amplifications in the resistant lines, and 41 non-silent single-nucleotide changes in 38 genes. Although none of the mutations was both common to all resistant clones, and wild-type in a revertant clone, mutations in SUMO were found in all mutants. Affinity purification with a benzoxaborole column yielded 14 proteins that bound specifically, but since none of these was affected in the resistant mutants, there was no clear indication as to which might be relevant to benzoxaborole action. Recent results have, in contrast, revealed that some anti-trypanosomal benzoxaboroles act as pro-drugs. Aminomethyl phenoxyl benzoxaboroles such as AN3056 are activated to aldehydes by a serum enzyme, then further modified intracellularly by a parasite aldehyde dehydrogenase to an active carboxylic acid form [26]. AN11736 is a slow-acting compound, although its EC_50_ is much lower than most of the benzoxaboroles. Selected resistance to this compound is associated with the loss of a serine peptidase that processes AN11736 to an active carboxylate derivative (Giordani *et al*., manuscript in preparation).

Various oxaboroles are being considered for treatment of apicomplexan parasites, and the compound AN13762 (Fig 1) is in development [27, 28]. So far, the only antimalarial benzoxaborole with a known mode of action is AN3661 (Fig 1). AN3661-resistant *Plasmodium* and *Toxoplasma* had mutations in the cleavage and polyadenylation factor CPSF3 [29, 30], which is responsible for cleavage of the mRNA precursor at the 3’ end. *In-silico* molecular docking of AN3661 into the predicted protein structure suggested that the boron atom occupies the position of the cleavage site phosphate of the mRNA substrate, with one hydroxyl group interacting with the zinc atom in the catalytic site. This is consistent with other benzoxaboroles which bind bimetal centers of beta-lactamase and phosphodiesterase-4 [28]. Introduction of some of the mutations - which were all in or near the active site - into susceptible parasites resulted in compound resistance. These results, combined with the loss of transcripts for three trophozolite-expressed genes in treated parasites, suggest that AN3661 inhibits mRNA polyadenylation through its interaction with CPSF3 [29, 30].

In trypanosomes, and indeed all *Euglenozoa* (the Order that includes Kinetoplastids), transcription of mRNAs is polycistronic. Individual mRNA 5’-ends are created co-transcriptionally by *trans* splicing of a 39nt leader sequence (*SL*) [31-33]. *Trans* splicing is spatially and mechanistically coupled to polyadenylation of the preceding mRNA. Polyadenylation sites are dictated by the positions of *trans*-splicing sites [34-37], RNAi-mediated depletion of polyadenylation factors inhibits trans splicing [38, 39], and disruption of splicing stops polyadenylation [34-37]. The spliced leader precursor RNA (*SLRNA*) is 140 nt long and is synthesised by RNA polymerase II [40] from approximately 200 tandemly repeated genes [41]. Unlike protein-coding genes, each *SLRNA* gene has its own promoter [42-44]. The *SLRNA* cap and the following four nucleotides are methylated [45-48]. Inhibition of *SLRNA* methylation using S-adenosyl-L-homocysteine or Sinefungin prevents splicing [49, 50]. This results in loss of mRNA from the cells by the normal mechanisms of mRNA turnover [51].

We here describe the results of a multi-pronged approach to discover molecular targets of several anti-trypanosomal benzoxaboroles. Our work concentrated on AN7973, which is orally active and was the DNDi back-up for SCYX-7158/AN5568 for the treatment of human African trypanosomosis.

## Results

### Anti-trypanosomal activities of AN7973

AN7973 was selected as a candidate veterinary drug from a range of 7-carboxamido-benzoxaboroles structurally similar to SCYX7158. The selection of AN7973 was based on its *in vitro* potency against *T. congolense* (Fig 1, S1 Table) and its ability to cure 5/5 *T. congolense*-infected mice with a single 10mg/kg i.p. dose, combined with *in vitro* ADME data. A proof of concept trial was then conducted using AN7973 in experimental goat models of infection against *T. congolense* and *T. vivax*. All *T. congolense*-infected goats (4/4 and 3/3) were effectively cured when AN7973 was administered as either two intramuscular injections of 10 mg/kg (given 24 hours apart) or as a single bolus dose injection of 10 mg/kg, respectively. Similarly, all *T. vivax*-infected goats (4/4) were effectively cured when AN7973 was administered as two intramuscular injections of 10 mg/kg (given 24 hours apart). However, a single bolus dose injection of 10 mg/kg was not sufficient to cure *T. vivax*-infected goats (0/4).

Encouraging cattle pharmacokinetic data following intravenous and intramuscular administration, and a good preliminary safety profile, prompted a cattle efficacy study of AN7973. The *T. vivax* and *T. congolense* isolates tested were both resistant to maximum dosages of diminazene (7 mg/kg) and isometamidium (1 mg/kg). A single 10 mg/kg intramuscular injection of AN7973 cured 3/3 cattle of *T. congolense* infection. Unfortunately, however, two 10 mg/kg injections of AN7973 cured only one out of two *T. vivax* infections, and a single injection failed to cure 3 animals. This meant that AN7973 would be inappropriate for field use [3]. The reduced efficacy of AN7973 against *T. vivax* might have been a consequence of weaker intrinsic potency (Fig 1, S1 Table). The *ex vivo* EC_50_ against *T. vivax* was 215 nM, as against an *in vitro* EC_50_ of 84 nM for *T. congolense* (S1 Table). The latter value is similar to the results for *T. brucei* (20-80 nM, S1 Table).

The lack of single-dose cattle efficacy at 10mg/kg i.m. against *T. vivax* precluded development of AN7973 as a commercially viable treatment against cattle trypanosomosis, but it could still have potential for diseases caused by other salivarian trypanosomes.

### Trypanosomes that are slightly resistant to AN7973 show many genome changes

One of the most direct strategies to identify the targets of antimicrobial drugs is selection of resistant mutants. We applied AN7973 selection to a library of *T. brucei* each of which over-expresses a different trypanosome protein or protein fragment [52]. When we grew this library in the presence of anti-trypanosome compounds with known targets, trypanosomes that ectopically expressed those targets were selected [53]. Upon growth of the over-expression library in the presence of increasing amounts of AN7973 we recovered two cell populations that grew in 65 nM and 80 nM AN7973. The over-expressed transgenes in these cells (Tb927.11.15930 and Tb927.8.6010) were however not able to confer resistance in newly transfected cells. We concluded that the very moderate resistance of the original cell lines must therefore have been due to mutations elsewhere in the genome.

To select more resistant parasites, we grew the two lines for four months in increasing amounts of AN7973, but the resulting cloned populations had EC_50_s of only 1.6x (75nM, clone 80C5 in the tables) and 2.1x (90 nM, clone 80D4 in the tables) relative to wild-type (45 nM). We subjected DNA from these clones, the clones from which they were derived (80C and 80D), and the original cells, to shotgun sequencing. We looked for mutations that affected protein sequences, and were found in more than one resistant line. S2 Table shows the changes in genes that are not part of multi-gene families ("unique" genes), while S2 Table and S3 Table detail all results. Nearly all of the changes were in multi-gene families so were unlikely to be significant. As previously observed for trypanosomes that were mildly resistant to AN2965 [25] we found amplification of numerous DNA regions, especially in the lines that had undergone more prolonged selection. For example, nearly 300 unique genes had increased copy numbers in both clone 80C5 and 80D4. The large number of genes affected made it impossible to pinpoint any particular pathway as being relevant to resistance (S2 Table).

### Metabolic effects of AN7973

To investigate the modes of action of AN7973 further we treated the original and the partially resistant lines with 10xEC_50_ (S1 Table) for 5h and characterised the metabolomes. There were no differences between the cell lines that gave any clue to the resistance mechanism, and no changes after treatment that indicated inhibition of one particular pathway. However, we did see at least 2-fold increases in methyl thioadenosine (MTA), methyl lysine, dimethyl lysine, trimethyl lysine, acetyl lysine, and S-adenosylmethionine (SAM) (Fig 1; S5 Table). SAM is a methyl donor and it has been shown that the methyl groups on the methylated lysines are derived from methionine (Steketee *et al.*, manuscript submitted). We had previously seen similar changes in these particular metabolites after treatment of trypanosomes with SCYX-7158 (AN5568, the sleeping sickness drug candidate) (Steketee *et al.*, manuscript submitted). It therefore seemed worthwhile to find out whether other anti-trypanosomal benzoxaboroles had similar effects.

We started with a panel of 30 compounds. In initial tests, in which trypanosomes were incubated with the compounds for 48h, 24 of the compounds showed sub-micromolar EC_50_s against bloodstream form *T. brucei*, (S1 Fig, S1 Table). The exceptions included AN3661, the malaria lead compound that targets CPSF3 [29, 30], which had EC50s for trypanosomes in the low micromolar range. Results for compounds that were selected for further work are shown in Fig 1. Some compounds acted within 6-8h while others only affected growth after 24h (S1 Fig, S2 Fig, S1 Table, S6 Table); the latter included the veterinary drug candidate AN11736, which requires processing for full activity (Giordani *et al*., manuscript in preparation). For *T. brucei*, EC_50_s were generally lowest for cells cultured in minimal media, and highest for procyclic forms (Fig 1). *In vitro* testing of ten compounds against *T. congolense* gave EC_50_s that were usually similar to those for *T. brucei*. The same was true for *ex vivo* tested *T. vivax*, although this species was clearly more susceptible than *T. brucei* or *T. congolense* to AN5568, but relatively resistant to AN3056 (Fig 1, S1 Table).

Treatment of *T. brucei* with 16 of the 30 compounds caused a statistically significant increase in SAM, and 15 of those compounds also caused an increase in MTA; results for selected compounds are in Fig 2 and all are included in S6 Table. There were no discernable structure-activity relationships for EC_50_, time to kill, or the SAM/MTA effect, but most of the oxaboroles that induced a SAM/MTA pattern were faster killing.

**Fig. 2.**
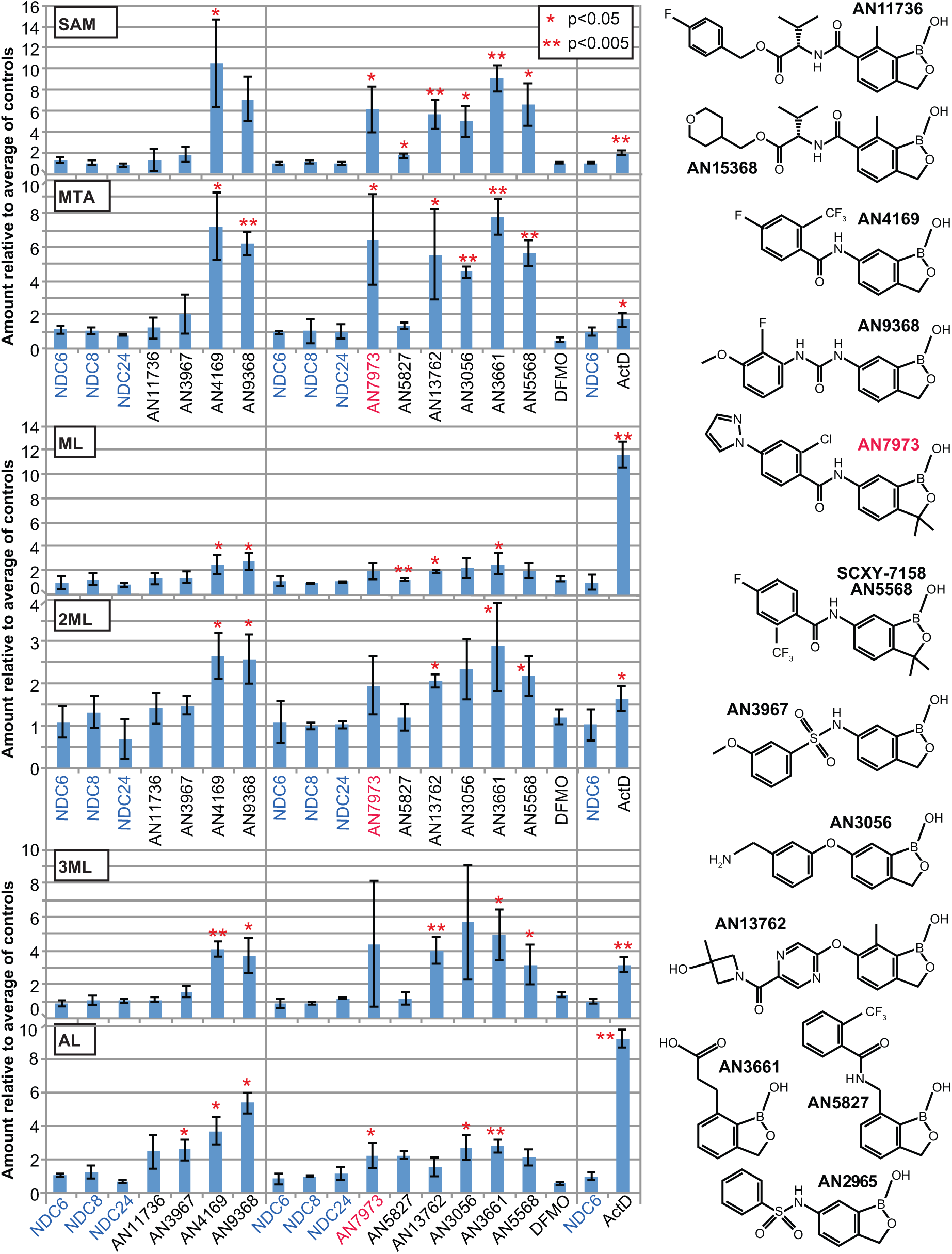
Effects of benzoxaboroles on the metabolome. Fold increases of selected metabolites for selected compounds. Structures are shown on the right. The numerical data are in S6 Table. SAM: S-adenosylmethionine; MTA: methylthioadenosine; ML: methyllysine; 2ML: dimethyllysine; 3ML: trimethyllysine; AL: acetyllysine.

### AN7973 inhibits protein synthesis

Since both resistance selection and metabolomics failed to reveal possible targets of AN7973, we reverted to more classical approaches. First we examined morphological effects. Jones et al. saw accumulation of parasites in the G2 phase of the cell cycle after AN2965 treatment [25]. Treatment of trypanosomes with AN7973 with 10x EC_50_ for 7h caused growth arrest (Fig 3A), but no obvious changes in parasite morphology or motility and no significant changes in the proportions of cells in different stages of the cell cycle (Fig 3B).

**Fig 3.**
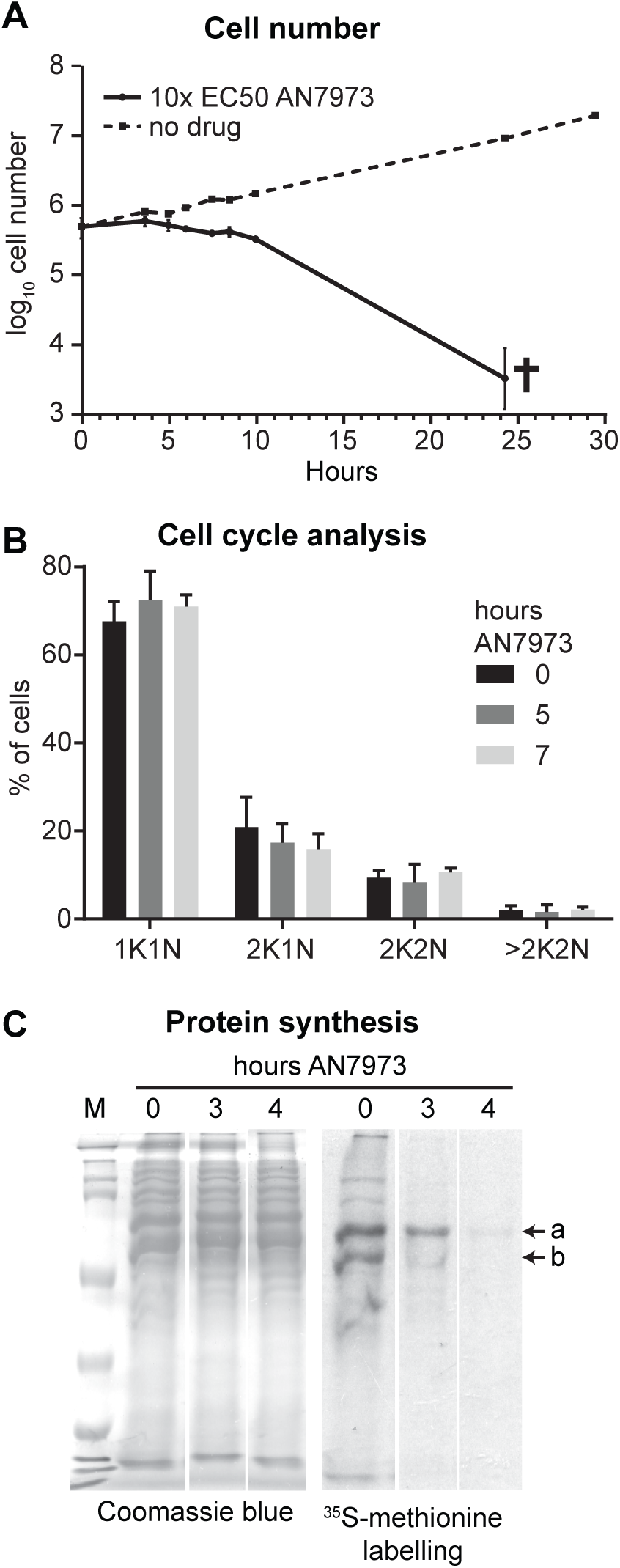
Effects of AN7973 on the cell cycle and protein synthesis A) Cumulative growth curve for bloodstream-form trypanosomes incubated with and without AN7973. Results are arithmetic mean ± standard deviation for three replicates. B) Cell cycle analysis by microscopy. The percentages of cells with different numbers of nuclei and kinetoplasts are shown. C) Effect on protein synthesis. 5×10^6^ Cells were incubated with AN7973 (10× EC_50_) for various times, then (after washing and 15-min pre-incubation) with [^35^S]-methionine for 30 min. Proteins were separated by SDS-PAGE and radioactive incorporation was assessed by autoradiography. The Coomassie-stained gel is shown as a loading control. Two prominent protein bands showing different effects of AN7973 are indicated as “a” and “b”. All lanes are from the same gel and exposure; the gaps between the lanes are present because identical results for two partially resistant lines (also treated at 10x EC_50_) have been deleted.

Amino-acyl tRNA synthetase inhibition is a known mode of action of several benzoxaboroles, and inhibition of tRNA synthetase should result in cessation of protein synthesis. We therefore measured this in AN7973-treated trypanosomes by pulse labelling with [^35^S]-methionine followed by denaturing polyacrylamide gel electrophoresis and autoradiography. Inhibition of protein synthesis (Fig. 3C) was clear, but some prominent protein bands were affected more than others (compare bands labeled a and b). The kinetics of the inhibition, combined with apparent selectivity for particular proteins, suggested that protein synthesis was not a primary target of AN7973, but might be inhibited in a secondary fashion.

### AN7973 inhibits mRNA processing

Loss of protein synthesis could be caused by loss of mRNA. For example, if mRNA synthesis were inhibited, the pattern in Fig. 3C would be explained if the mRNA encoding protein “b” were less stable than that encoding protein “a”. Loss of mRNA could be caused by inhibition of either RNA transcription or processing. To investigate this possible mechanism, we incubated cells with AN7973, prepared RNA, and hybridised Northern blots with a probe that detects the spliced leader *SL.* This probe detects all processed mRNAs, as well as the *SLRNA* that donates the *SL*. Incubation with AN7973 for 9h had little or no effect on the total amount of rRNA, as judged by methylene blue staining (Fig. 4A) but caused progressive loss of spliced mRNA (Fig. 4B). In contrast, the levels of *SLRNA* remained roughly constant (Fig. 4B), suggesting that RNA polymerase II transcription of *SLRNA* genes was not inhibited. We therefore suspected that AN7973 was inhibiting mRNA processing. To test this we re-hybridised the blot with a probe that detects beta-tubulin. The tubulin genes are arranged in alternating alpha-beta tandem repeats, and splicing inhibition through heat shock [54] or Sinefungin treatment [49] leads to accumulation of partially processed beta-alpha dimers and multimers. Partially processed tubulin mRNAs were indeed detected within an hour of AN7973 application (Fig. 4C), indicating that transcription continued but processing was impaired. At later time points, the tubulin mRNAs disappeared, suggesting that the failure in processing was complete. (Most measurements suggest that the tubulin mRNAs have half-lives of about half an hour [55].)

**Fig 4.**
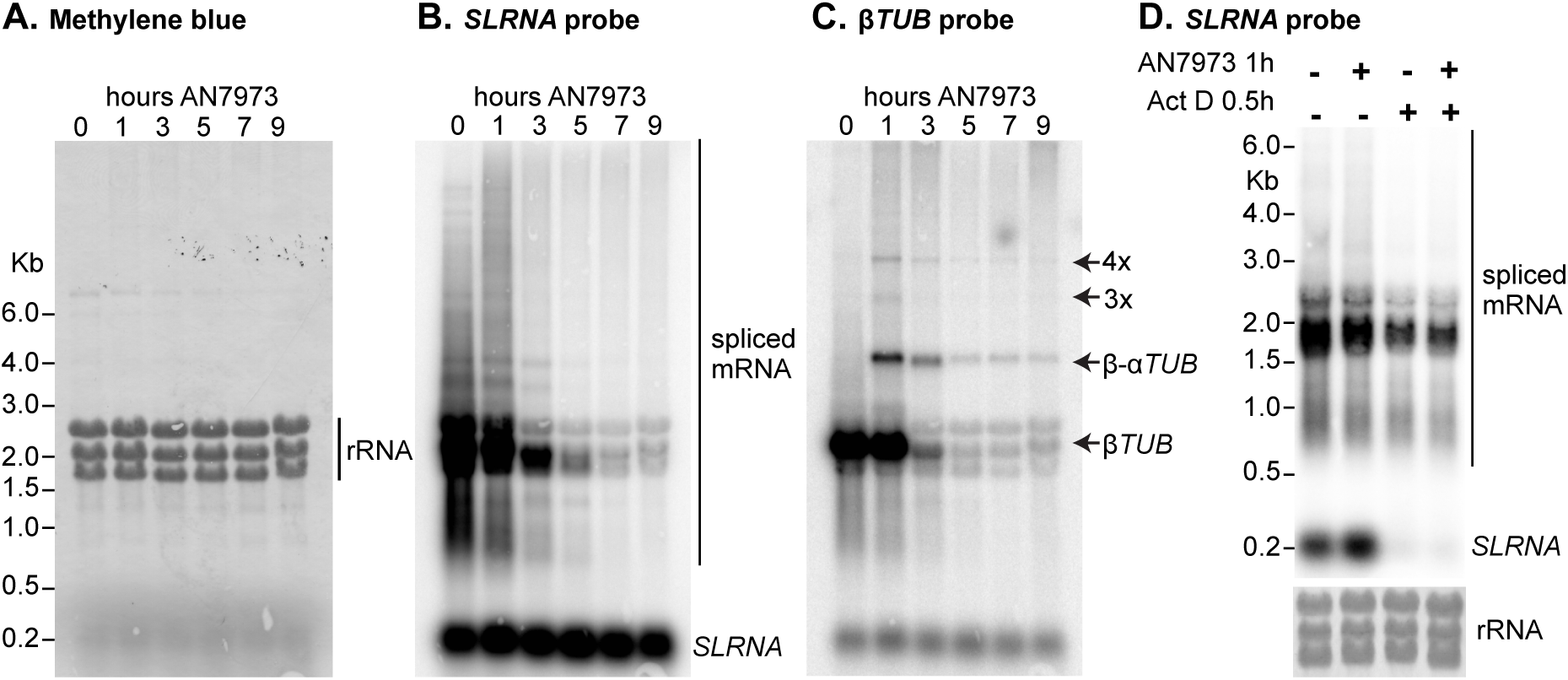
AN7973 affects mRNA processing A) Trypanosomes were incubated with AN7973 for different times (0-9 hours). RNA was subjected to denaturing agarose gel electrophoresis, blotted, and detected using radioactive probes. This panel shows Methylene blue-stained gel showing equal loading of total RNA. The three prominent bands are rRNA. B) As (A), blot probed with a [^32^P]-end-labelled probe complementary to the spliced leader. The signals representing the ~140nt *SLRNA* and mature mRNAs are indicated. The cluster of bands around 2 kb probably represent the abundant mRNAs encoding the Variant Surface Glycoprotein, alpha and beta tubulin (both ~1.9 kb including 100nt poly(A)), and EF-1 alpha (1.8 kb) [79]. C) The blot from (B) was stripped then incubated with a beta-tubulin probe. Shadows from the *SL* probe remain. The positions of monomer, dimer and multimers from the tubulin repeats are indicated. D) The effect of AN7973 is not due to inhibition of *SLRNA* synthesis. Trypanosomes were treated either with AN7973 for 1h, or with Actinomycin D for 30 min, or with AN7973 followed by Actinomycin D, as indicated above the lanes. The lower panel is the rRNA loading control.

As noted above, after 1h AN7973 treatment, the level of *SLRNA* was either unchanged or slightly increased. RNAi targeting splicing or polyadenylation factors has similar effects [39, 56, 57]. For example, RNAi that targeted polyadenylation factors caused 2-4 fold increases in *SLRNA* [39]. The absence of a massive increase in *SLRNA* can be explained by a balance between continued *SLRNA* transcription and degradation of the unused *SLRNA*. After Actinomycin D treatment to inhibit transcription, the pre-existing *SLRNA* decreases even though no new precursors are available for splicing [58]. Fig 4D compares the effects of AN7973 with those of Actinomycin D treatment. As expected, 30 min Actinomycin D resulted in *SLRNA* loss; and if Actinomycin D was added after the AN7973 treatment, the *SLRNA* again disappeared. All results were therefore consistent with AN7973 inhibiting splicing without affecting *SLRNA* transcription.

### mRNA processing inhibition by AN7973 is not due to loss of cap methylation

The only small molecules currently known to inhibit kinetoplastid mRNA processing *in vivo* are S-adenosyl methionine analogues such as Sinefungin, which are general methylation inhibitors [59]. They inhibit *trans* splicing by preventing spliced leader methylation [49]. To find out whether AN7973 had the same mechanism, we used primer extension to measure the amounts of *SLRNA* and of the 2’-5’ branched “Y-structure” splicing product (Fig. 5A). cDNA synthesis was primed by a 5’-labelled oligonucleotide that is complementary to a region towards the 3’ end of the *SLRNA* (black arrow in Fig. 5B). The products from full-length *SLRNA* form a small ladder, because 5’ methylation partially blocks reverse transcriptase (Fig 5B). In addition, the Y-structure trans splicing intermediate gives a product of 87nt (Fig. 5B). These products are seen in Fig 5C, lane 1. As expected, incubation with the methylation inhibitor Sinefungin for 30 min abolished the multiple bands caused by cap methylation, with cDNA synthesis extending cleanly to the 5’ end of the *SLRNA*; at the same time, the signal from the Y structure was decreased (Fig 5C, lane 4, arrowhead). The standard concentration of Sinefungin used, 2 µg/ml, is sufficient to prevent cap methylation within about 10 min, and is more than 4000 times the EC_50_. Unlike Sinefungin, AN7973 (10x EC_50_) had no effect on the pattern of bands from the *SLRNA*, showing that cap methylation was not affected. Nevertheless, Y-structure formation was clearly decreased, confirming splicing inhibition (Fig. 5C, lanes 2 and 3). Quantitation of replicate independent experiments showed that a 1-h incubation with AN7973 inhibited *trans* splicing to the same extent as a 30-min incubation with Sinefungin (Fig 5D), and that after 30 min, the effect was already significant (p<0.0005) (Fig 5E). In contrast, diminazene aceturate, suramin, eflornithine and pentamidine had no significant effect in the same assay (Fig. 6), showing that splicing inhibition was not simply a side-effect of growth inhibition.

**Fig 5.**
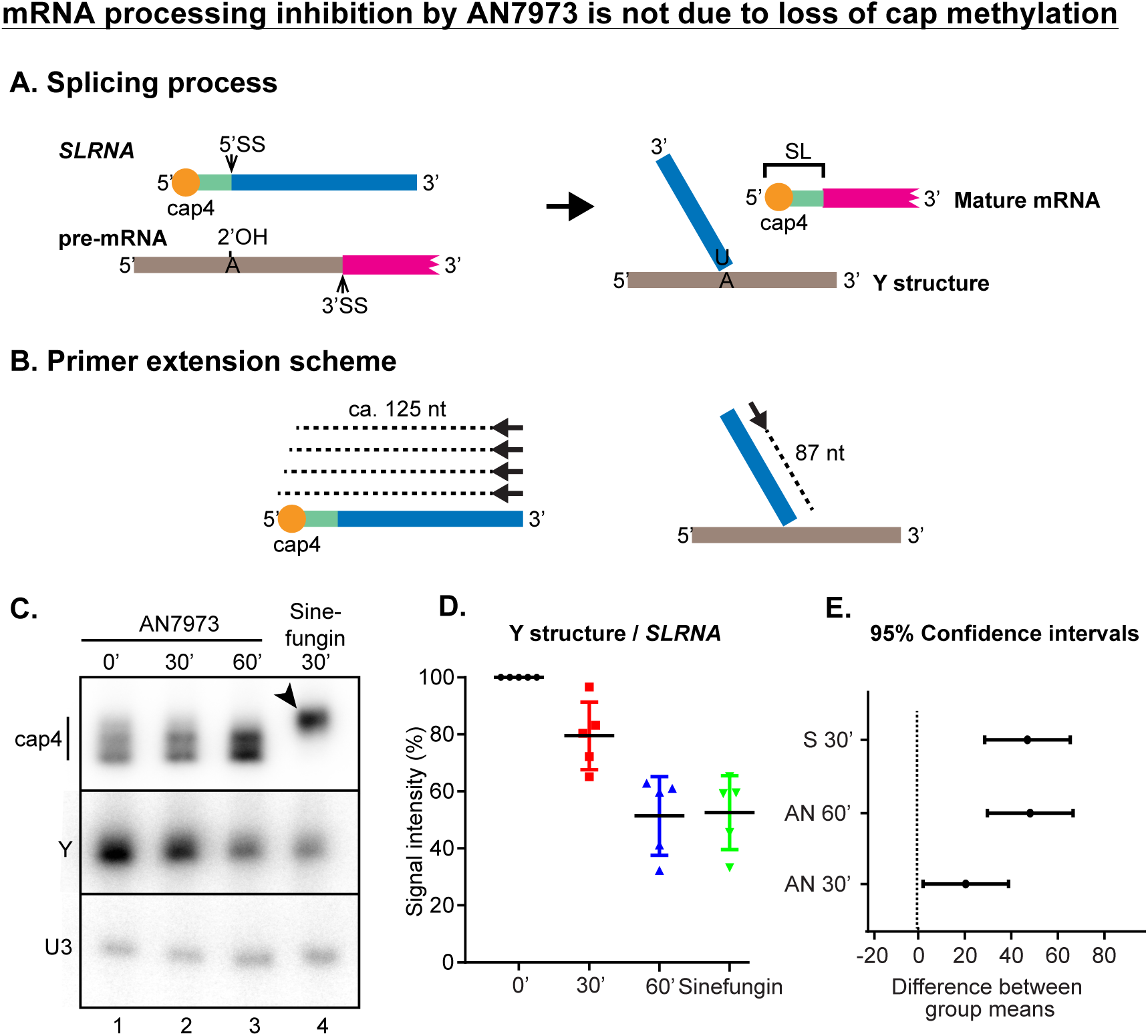
Effect of AN7973 on *trans*-splicing. A) Schematic representation of the splicing reaction. *SLRNA* and Y structure during primer extension experiments. *SLRNA* is represented with various colours; red is the cap4 region, green is the spliced leader (*SL*), blue is the *SL* intron. The pre-mRNA is shown with a grey intergenic region and a magenta portion that represents the 5’ end of a mature mRNA. The 5’ and 3’ ends of each RNA are indicated. During *trans* splicing, the SL intron forms a branched “Y” structure with the 2’ hydroxyl of an adenosine located 5’ to the polypyrimidine tract that is recognised by the splicing machinery. Meanwhile the *SL* is *trans* spliced to the 5’-end of the mRNA. B) Schematic representation of the primer extension assay. The 5’-end labelled oligonucleotide primer hybridises towards the 3’ end of the *SLRNA*. Reverse transcription on the intact *SLRNA* extends (dashed line) towards the 5’ end but terminates upon encountering the methylated residues of cap4. In the Y structure, the primer extends (dashed line) until the branch point, giving an 87nt product. C) Typical result from a primer extension experiment illustrated in (B). A primer complementary to the U3 snRNA was used as a control. Lane 1 (0’) shows the result from trypanosomes that were treated with DMSO only. Lanes 2 and 3 show the results from trypanosomes treated with AN7973 for 30 min and 60 min, and lane 4 is the result for 30 min Sinefungin. The arrowhead shows the primer extension product from unmethylated *SLRNA*. D) Quantification from four independent experiments; the ratio between Y structure and the *SLRNA* signal is shown. After only 30 min incubation with AN7973, the difference is significant using a one-way ANOVA E) 95% confidence intervals of mean differences between Y structure quantifications, calculated using Dunnet’s multiple comparisons test. In each case the calculation is done for the comparison between no treatment and the treatment indicated. The higher the difference, the lower the chance that the values were not significantly different. A value of 0 indicates a 5% chance that differences were not significant.

**Fig. 6.**
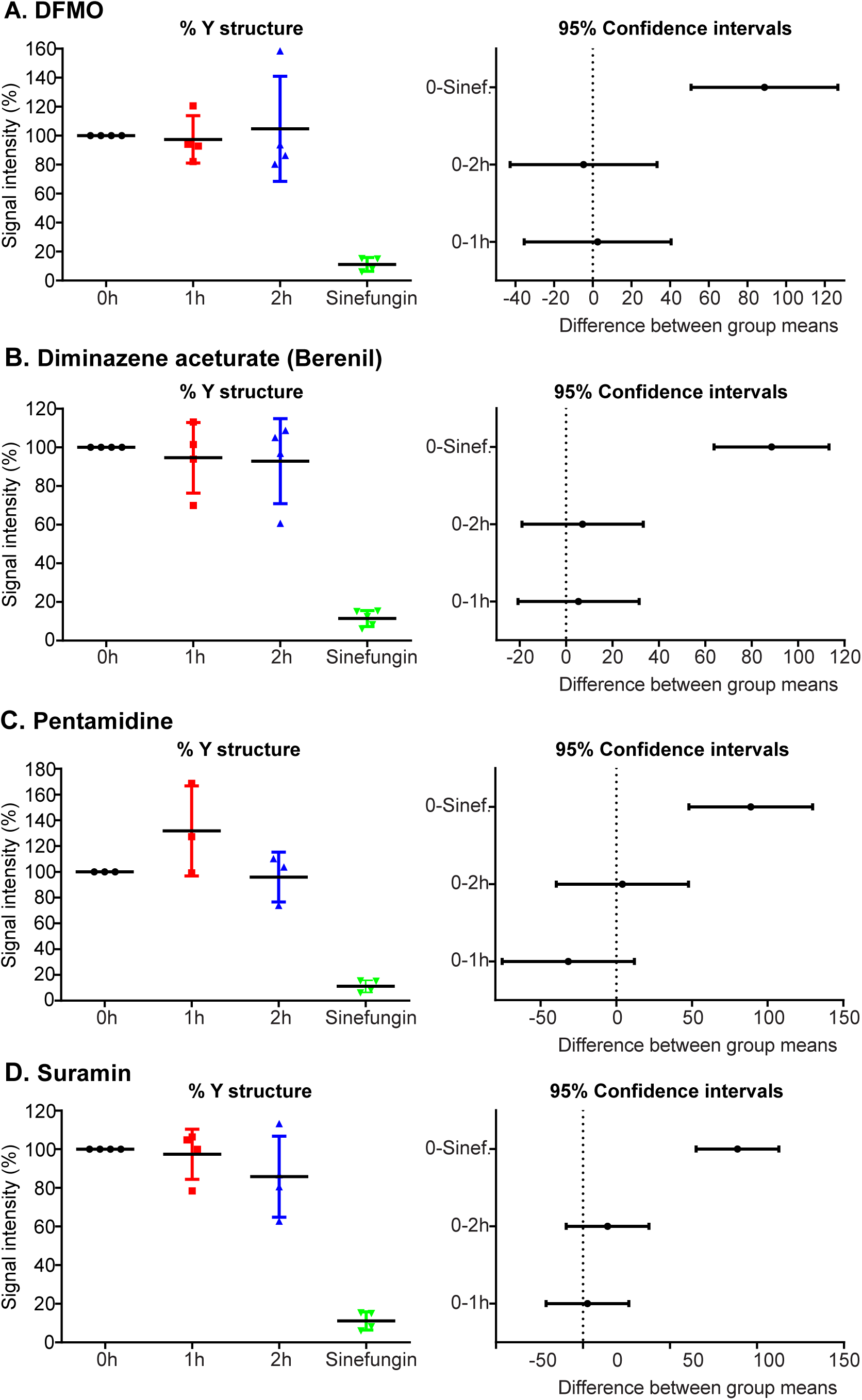
Specificity of *trans* splicing inhibition Trypanosomes were incubated with 10x EC50 of various known anti-trypanosomal drugs for 1h or 2h, then splicing was assayed as in Fig. 4. The % Y structure (relative to U3) is on the left and the 95% confidence intervals on the right.

### Relationship between splicing inhibition and SAM/MTA accumulation

At this point, we wondered whether there was any link between the metabolome changes caused by AN7973 treatment and splicing inhibition. A possible reason for the increase in S-adenosyl methionine could be decreased requirements for cap methylation. Transcription and processing of the *SLRNA* genes absorbs substantial cellular resources, since the cell needs to make at least 10000 *SLRNA*s per hour [55]. It was possible that after 5-6h, there might have been some feedback inhibition of transcription that led to a decreased methylation requirement. An alternative explanation is that loss of mRNA production leads to loss of unstable mRNAs. If the protein encoded by an unstable mRNA has a high turnover rate, then that protein will disappear; and if that protein is a metabolic enzyme, its substrates will accumulate. This too could have led to metabolic changes as we noted.

To investigate the possible link between splicing inhibition and methylation we measured their effects of various compounds on splicing. In addition to a selection of compounds that had been metabolically analysed, we included AN2965, the compound whose mode of action had been had been investigated previously. Five compounds out of six that caused increased SAM/MTA showed a decrease of 40% or more in the Y structure, similar to the effects seen with Sinefungin and AN7973 (Fig 7, S3 Fig A). The processing inhibitors included the antimalarial candidate AN13762 (compound 46 in [28]); the EC_50_ of this compound for trypanosomes in the 3-day assay was 4 times higher than that for *P. falciparum*. The exceptional compound was AN3056, which had a SAM/MTA effect but gave only 20% Y-structure inhibition (S3 Fig A). Of the compounds that did not give SAM/MTA effects, AN11736 had only a minimal effect on splicing. Since results from AN3967 and AN5827 were too variable to interpret, we re-examined them, as well as AN11736, in a second experimental series. This time, inhibition was observed although not as strong as for AN7973 (Fig 7). More reproducible, or stronger, inhibition might be obtained after a longer incubation but it is also possible that the effects on splicing from these compounds were secondary to other stresses [54].

**Fig. 7.**
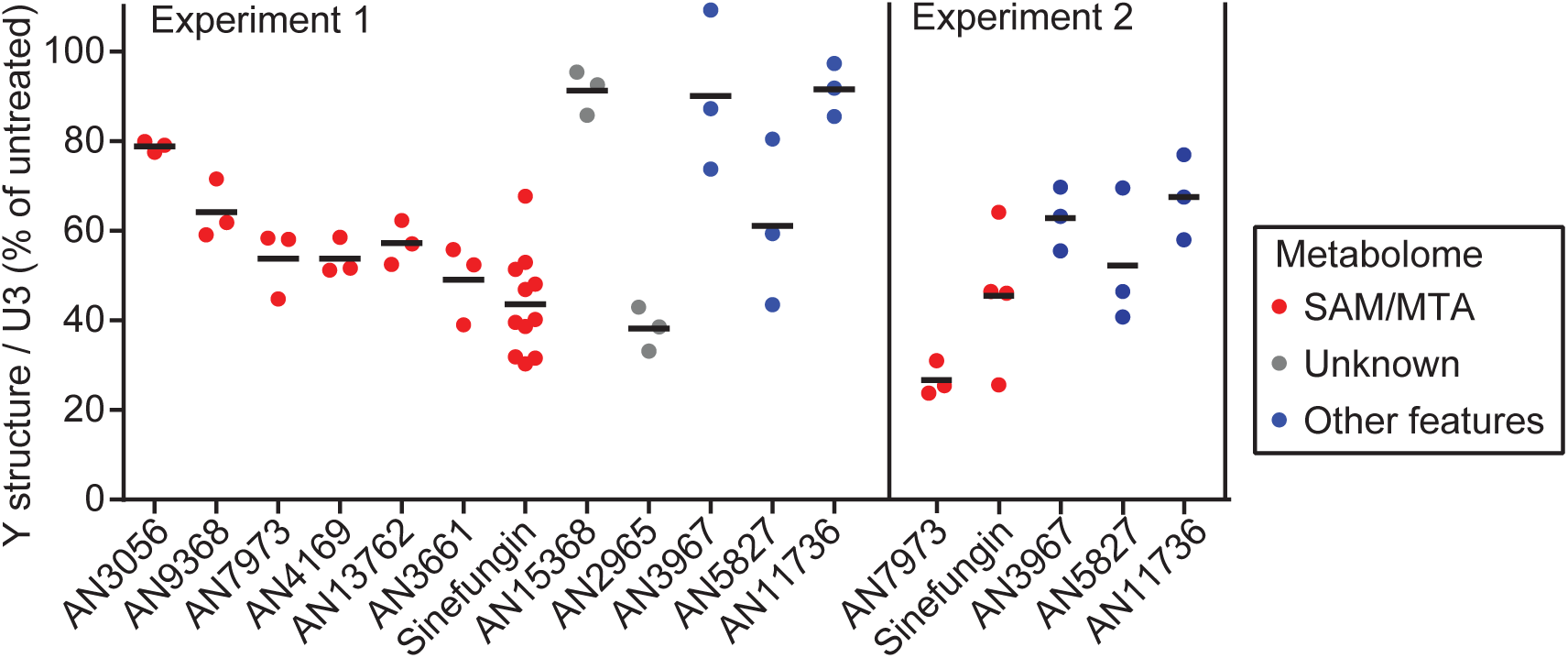
Effects of various benzoxaboroles on *trans* splicing Trypanosomes were incubated with 10x EC50 of various compoundss for 1h or 2h, then splicing was assayed as in Fig. 3. The % Y structure (relative to U3 as control) is shown. Results are shown for one experiment (experiment 1) with a full compound set and a second experiment for selected compounds.

We also directly addressed the effect of mRNA synthesis inhibition on the metabolome, by incubating with the standard concentration of Actinomycin D (10 µg/ml) for 6h. There were large increases in methylated amino acids but the increases in SAM and MTA were lower than those seen with many of the oxaboroles (Fig 2, S6 Table). There was also a weak correlation with the effects of AN5568 (S3 Fig B). We concluded that loss of mRNA might at least partially explain some of the effects of AN7973 on the metabolome.

### In vitro splicing

As an alternative way to find out whether the effect of AN7973 on mRNA processing was direct or indirect, we measured *trans* splicing in permeabilised procyclic-form trypanosomes [50]. (Equivalent assays are not established for bloodstream forms.) as for most benzoxaboroles that have been tested (S1 Fig, S1 Table) the EC_50_ of AN7973 for procyclic forms was 5-10 times higher than that for bloodstream forms. Treatment with 10x EC_50_ clearly inhibited splicing in procyclic forms, with 70% loss of Y structure after 2h (S4 Fig). To test splicing, procyclic trypanosomes were permeablised with lysolecithin, pre-incubated with AN7973 or DMSO, then transcription was allowed to proceed for 10 min in the presence of [alpha-^32^P]-UTP [60]. RNA was separated on 7% polyacrylamide-urea gels and visualized by autoradiography. Under these conditions, the *SL* intron is visible as a ~100nt species, which disappears if incubation is continued for a further 20 min [60]; it is also not made if cap methylation is inhibited by S-adenosyl homocysteine [50].

We do not know the intracellular concentration of AN7973, and in the *in vitro* transcription reaction the density of permeabilised parasites is 1000 times higher than in the experiments with cultures. For the in vitro assays we therefore chose to use a concentration of 100 µM (500x the EC_50_), which gives a AN7973:parasite ratio that is equivalent to the ratio at 5x EC_50_. This treatment reproducibly prevented formation of the Y structure without preventing transcription of *SLRNA* or smaller RNAs (Fig 8). An additional band (arrowhead), which appeared in the presence of AN7973 and was slightly shorter than *SLRNA*, might be a 3’ degradation product that was previously described [50]. AN7973 also reproducibly inhibited labeling of RNAs longer than 500nt (indicated by a question mark). These RNAs are thought to include mRNAs and rRNA [61], and the inhibition could either be a consequence of *trans* splicing inhibition or another effect of AN7973. In future it would be interesting to repeat these studies at a variety of concentrations and with other benzoxaboroles.

**Fig. 8.**
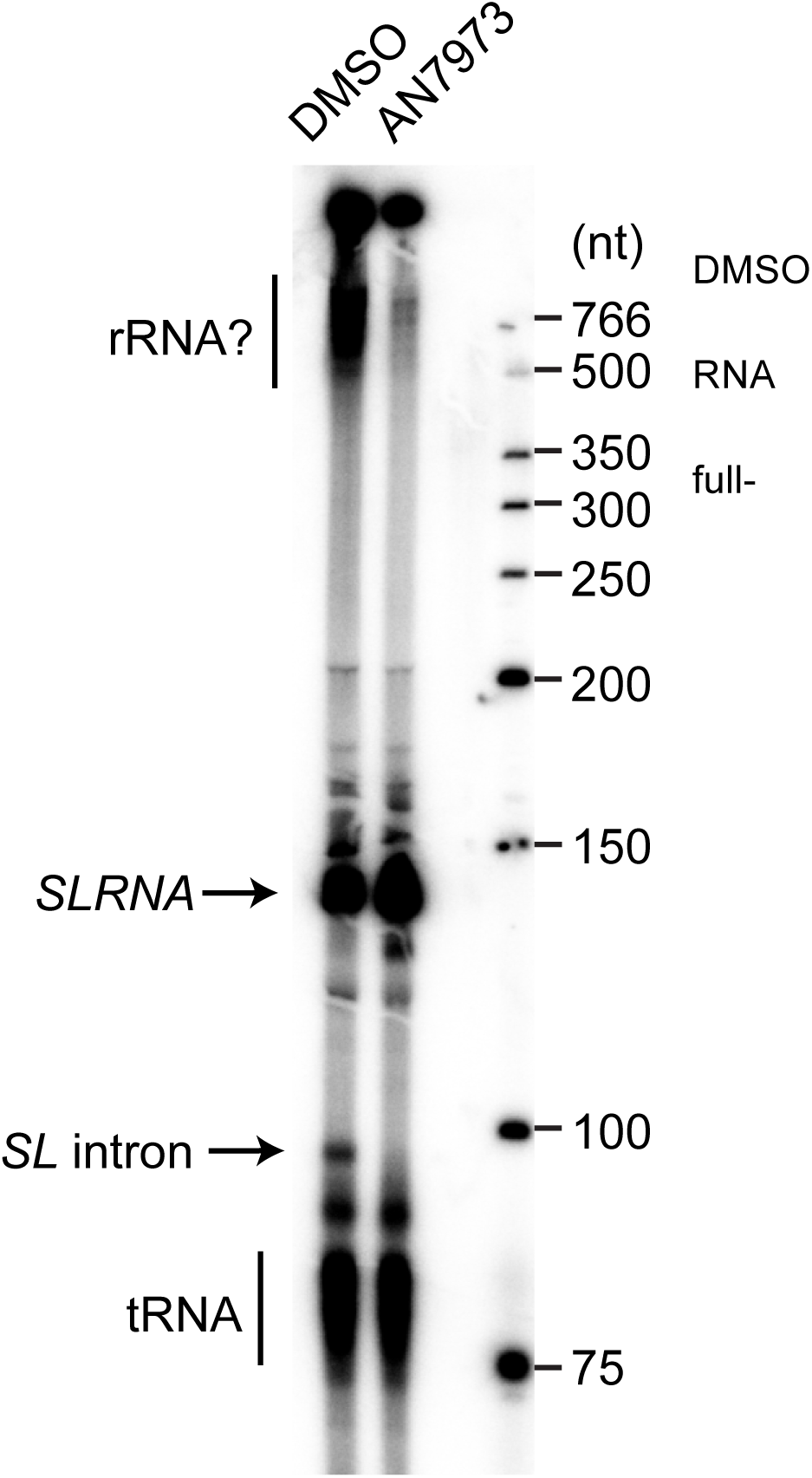
*In vitro* splicing assay. Permeablised procyclic-form trypanosomes (2.5 × 10^8^ / reaction) were incubated for 2 min in the presence of or AN7973, and then for 10 min in the presence of alpha-32P-UTP. Newly-made was analysed by denaturing gel electrophoresis and autoradiography. The length SLRNA (~140nt) and the debranched SLRNA intron (~100 nt) are indicated.

## Benzoxaborole treatment induces formation of stress granules

When trypanosomes are stressed by heat shock or starvation, their mRNAs accumulate in RNA-protein particles called stress granules [62]. The granules contain the helicase DHH1 and are distributed throughout the cytoplasm. After 30 min Sinefungin treatment, or 2h after electroporation of a morpholino oligonucleotide targeting the U2 snRNA, DHH1 is also found in granules, but they are arranged in a characteristic pattern surrounding the nuclear envelope [63]. The formation of these “nuclear periphery granules” depends on the presence of partially processed mRNAs [63]. The transient appearance of DHH1-containing nuclear periphery granules should therefore distinguish specific splicing inhibition from more non-specific stress responses. The disadvantage of this assay is that the effect of splicing inhibition on DHH1 localization is time-dependent: upon more prolonged Sinefungin treatment some of the granules move away from the nucleus and become distributed throughout the cytosol, like other types of stress granule.

To analyse DHH1 localization after benzoxaborole treatment, we created bloodstream-form trypanosomes in which one *DHH1* open reading frame had been fused to a sequence encoding yellow fluorescent protein (YFP). We then examined the location of the YFP-DHH1 fusion protein (Fig 9; note that the YFP signal is coloured magenta). A few granules were seen in untreated cells, probably induced by washing and centrifugation (Fig 9A). After 30 min Sinefungin treatment, the nuclear periphery granule pattern was clear (Fig 8B). AN7973 also induced formation of perinuclear granules in a subset of cells (Fig 9C). We therefore, in three separate experimental series, treated the cells with different benzoxaboroles for 2h, and (after blinded renaming of the image files to prevent bias) quantitated DHH1 granule formation. Untreated cells formed fewer granules than drug-treated cells, and hardly ever showed granules around the nucleus (Fig 9A, D). Although results from three replicate treatments were variable (S5 Fig), most of the benzoxaboroles caused an increase in the numbers of granules, with perinuclear patterns in at least some cells (Fig 9D, S6 Fig, S7 Fig). There was strikingly little correlation between granule formation, Y structure inhibition and SAM/MTA accumulation (Fig 9D, S3 Fig C).

**Fig. 9.**
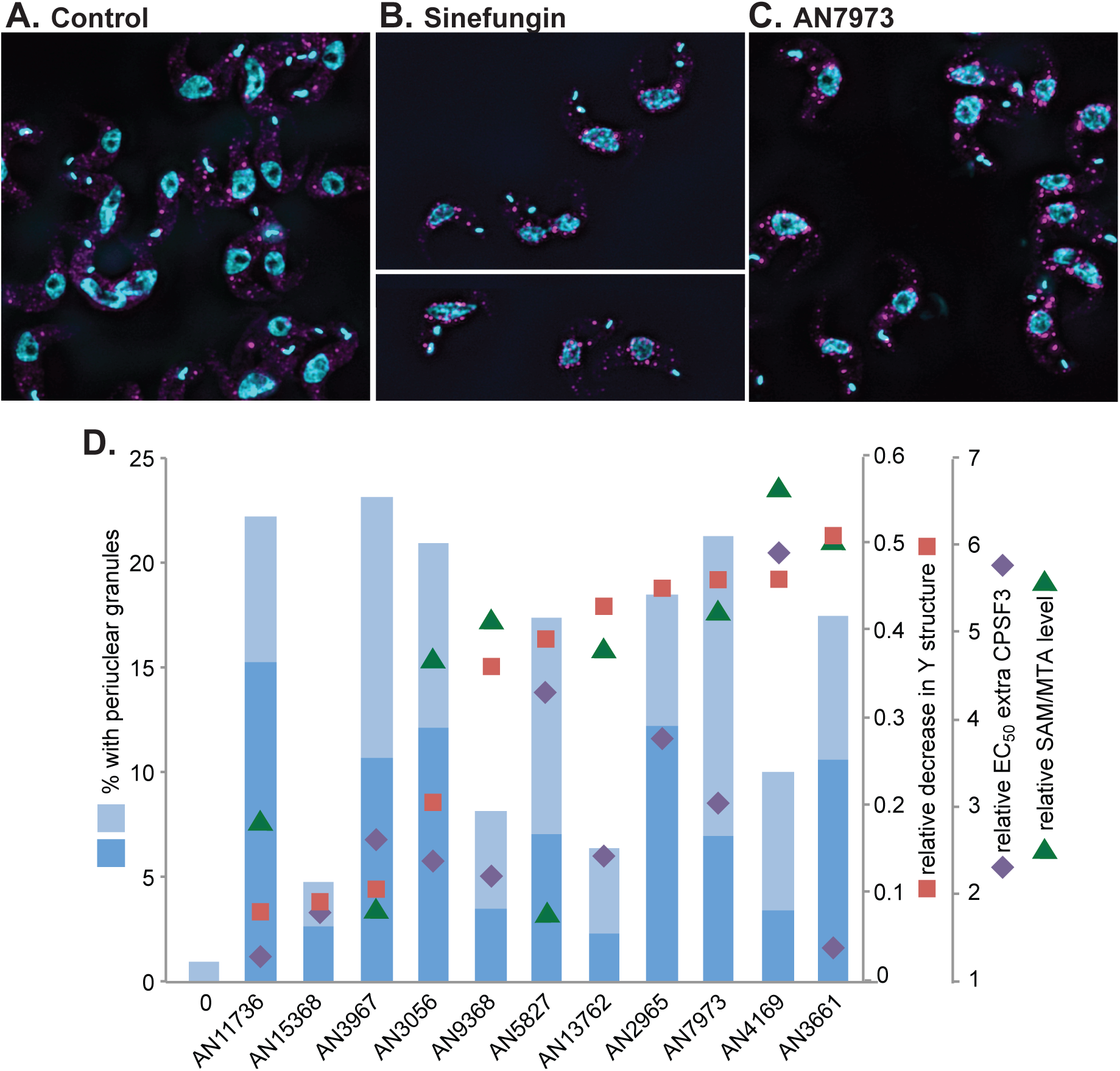
Effects of benzoxaborole treatment on formation of YFP-DHH1 granules Bloodstream-form trypanosomes with one *DHH1* gene tagged *in situ* were washed with PBS, fixed with formaldehyde, then allowed to adhere to glass slides before imaging to detect YFP (magenta) and DNA (cyan). Cells were either shown without pre-treatment (A) or after incubation with Sinefungin (1µg/ml 30 min) (B) or AN7973 (230 nM, 2h) (C). D) Results for all tested oxaboroles. The average percentages of cells with clear (dark blue) or possible (pale blue) peri-nuclear granules are shown. Results for the three replicate experiments are in S5 Fig. Results of other assays as indicated by symbols, as indicated on the Y-axes on the right-hand side.

### Is CPSF3 the target of AN7973?

To try to identify possible targets implicated in mRNA processing, we re-examined the genomes of our partially-resistant lines. No mutations of the U snRNAs were found. The only splicing factor gene with a missense mutation is annotated as “CRN/SYF3 part of splicing PRP19 complex, putative”, (Tb927.10.9660) which was found in the purified PRP19 complex [64]. The Sm complex forms the core of spliceosomal snRNPs, and our two cell lines that had been selected in AN7973 for several months also both had 1-2 extra copies of the genes encoding four out of the seven components: Sm-B, Sm-E, Sm-F and Sm-G. The other three genes were, however, not amplified. After selection of proteins on an oxaborole affinity column, Jones et al. found enrichment of RBSR1 (Tb927.9.6870), a protein with an SR-domain that is potentially involved in splicing, and of the U2 splicing auxiliary factor (U2AF35, Tb927.10.3200) [25]. In one of their AN2965 resistant lines, Jones *et al.* [25] observed a two-fold amplification of the gene encoding CPSF3 (Tb927.4.1340; also designated CPSF73). Our resistant lines had no changes in CPSF3, RBSR1 or U2AF35.

In trypanosomes, loss of polyadenylation stops splicing [38, 39], and in *P. falciparum* and *T. gondii*, mutations in CPSF3 gave resistance to AN3661 [29, 30]. CPSF3 therefore seemed the most promising candidate for follow-up. We first compared the *T. brucei* sequence with those of *Plasmodium* and humans, concentrating on the residues that were mutated in AN3661-resistant Apicomplexa. H36 and Y408 of the *P. falciparum* sequence were mutated in AN3661-resistant lines but were retained as H and Y the human and trypanosome sequences (S8 Fig). The remaining important *Plasmodium* residues, however, were already different in trypanosomes. Y252 (mutated to C for resistance) is N in *T. brucei*; T406 (mutated to I) is A; T409 (mutated to A) is C; and D470 (N) is already N in *T. brucei* (S5 Fig). These changes may explain why the IC50 of AN3661 for trypanosomes is high (S1 Fig). We attempted to make the equivalent of the Y408S mutation in trypanosomes, by homologous gene replacement, but were unable to obtain viable cells with the mutation. This might be a technical problem, but it is also possible that in the context of the trypanosome sequence, the Y408S mutation results in an unacceptable decrease in enzyme activity. If so, the result suggests either that the mutant CPSF3 has dominant-negative effects, or that CPSF3 is present in limiting amounts, so that even loss of one active gene copy is not compatible with growth.

As an alternative to mutation, we inducibly over-expressed RBSR1, U2AF35 and CPSF3 in bloodstream forms, as C-terminally myc-tagged versions. Expression of RBSR1-myc and U2AF35-myc did not affect the EC_50_ of AN7973. In contrast, expression of CPSF3-myc (S9 Fig A) caused at least 3-fold increases in the EC_50_s of four tested benzoxaboroles, including AN7973 (S9 Fig B, S7 Table). More modest (but statistically significant) increases were also seen for other benzoxaboroles (S9 Fig B, S7 Table) but the correlation with Y structure formation was weak (S3 Fig D). Over-expression of CPSF3-myc had little effect on the EC50 of AN3661, which is not surprising given the number of relevant amino-acid changes relative to the Apicomplexan enzyme; this suggests that AN3661, which was applied in micromolar concentrations, might be inhibiting splicing via a different target or indirectly. We suggest that the benzoxaboroles with 3x increased EC_50_s in cells with CPSF3-myc expression probably can bind to CPSF3. This would partially rescue the cells because the free intracellular compound concentration would be effectively reduced.

## Discussion

Benzoxaboroles are important drug candidates for both human and ruminant African trypanosomosis. Our results show that AN7973 inhibits mRNA processing in trypanosomes. AN7973 caused accumulation of DHH1-containing perinuclear granules that are typically found after splicing inhibition. Expression of additional CPSF3 increased the EC_50_ of AN7973 3-fold, suggesting that AN7973 can bind to CPSF3. These results suggest that mRNA processing is an important target of AN7973, which might operate through CPSF3 inhibition. AN7973 also caused metabolite changes indicative of disturbed methylation, similar to those observed for acoziborole.

When this work was started, AN7973 was under consideration as a candidate for treatment of cattle trypanosomosis, but it was later found to be relatively ineffective against *T. vivax*. In the available *T. vivax* genome sequence there is a gap in the CPSF3 open reading frame which includes Asn231 (*T. brucei* numbers), which is Tyr in Apicomplexa and humans, and mutated to Cys in resistant Apicomplexa. All of the other CPSF3 residues that are implicated in AN3661 resistance in *Apicomplexa* are conserved between *T. vivax*, *T. congolense* and *T. brucei* (S10 Fig).

It is not clear whether there is a mechanistic link between splicing inhibition and methylation defects (Table 1, Fig 8). One hypothesis we had was that the effects on methylation intermediates could be caused by decreased requirements for cap methylation. Transcription and processing of the *SLRNA* genes absorbs substantial cellular resources, since the cell needs to make at least 10000 *SLRNA*s per hour (one for every mRNA) [55]. The argument against this is that we found no evidence, either *in vivo* or *in vitro*, that *SLRNA* transcription, capping and methylation were affected by AN7973. Another hypothesis was that loss of mRNA production will lead to a selective reduction in the activities of enzymes that have relatively high mRNA and protein turnover rates, and that these enzymes are required for various aspects of methylation. This was partially supported by accumulation of methylated amino acids after Actinomycin D treatment (S6 Table). However whilst Actinomycin D did cause an increase in SAM it was less pronounced than with AN7973. This may mean that the changes in methylated lysine have different causes after the two treatments.

Our results with various other benzoxaboroles revealed no discernable correlation between trypanosome killing, granule formation, the metabolome, or splicing (Fig 8, Table 1). The diversity in structures of compounds that inhibited of trypanosome mRNA processing suggests to us that this may be an intrinsic property of the pharmacophore, limited perhaps by side-chains that block access or by requirements for compound accumulation. Nevertheless, some trends with respect to molecules with the same scaffolds can be noted. For example the two compounds AN15368 and AN11736 are both L-valinate amide benzoxaboroles [24] and both showed very little Y structure inhibition (Figs. 6 & 8). These two compounds also showed little change in EC50 with overexpression of CPSF3 (Fig 8) suggesting that the L-valinate amide benzoxaborole series may not act through inhibition of mRNA processing. AN7973 and AN4169, both of the carboxamide scaffold, also showed similar decreases in Y structure. An important caveat is that the uptakes and metabolisms of the various compounds are likely to differ. For example, after 2h incubation, AN3056 and the veterinary drug candidate AN11736 had less effect on processing in than did AN7973: but their action might be delayed, since both are subject to activation within the parasites. Evidence so far indicates that the formation of nuclear periphery granules containing DHH1 is caused by splicing inhibition [62]. However, we found no correlation between granule formation and inhibition of Y-structure formation. The absence of peri-nuclear granules after treatment of cells with benzoxaboroles that do cause splicing inhibition could be explained by the time-dependency of the assay. In contrast, the presence of peri-nuclear granules in the absence of detectable splicing inhibition suggests either slight differences in kinetics (granules seen before an increase in Y-structure?) or that this pattern can be produced by other mechanisms.

So far, selection of trypanosomes for resistance to with benzoxaboroles has met limited success. Substantial resistance was obtained only for compounds which require intracellular metabolic activation, and the mutations responsible were in the activating enzymes. For benzoxaboroles that are probably not metabolised, such as AN7973 and AN2965, only very limited resistance could be obtained. If these compounds had a single enzyme target, such as CPSF3, selection for resistance ought to have been relatively straightforward. The difficulty in obtaining mutants therefore suggests to us that anti-trypanosomal benzoxaboroles studied here may have several cellular targets. Nevertheless, inhibition of mRNA processing is so far the earliest effect that has been seen, and is likely to make a vital contribution to parasite killing. This raises the possibility that similar modes of action might be seen in oxaboroles under development against other Kinetoplastids, including *Leishmania* and *Trypanosoma cruzi*.

## Materials and Methods

### Compound efficacy studies for *T. congolense* and *T. vivax*

Testing of *T. congolense in vitro* (72 hrs) and *T. vivax ex vivo* (48 hrs) assays were done as described in [65]. Testing of *T. congolense* and *T. vivax in vivo* in mice was done as described in [24]. All *in vivo* mouse experiments were carried out in accordance with the strict regulations set out by the Swiss Federal Veterinary Office, under the ethical approval of the Canton of Basel City, under license number #2813.

The proof of concept efficacy studies in goats using AN7973 were conducted as previously described in [66], but established and modified for *T. congolense* and *T. vivax* models of infection. The trials took place from January to May 2013 within the Veterinary Faculty of the University of Las Palmas and the Agricultural farm (Granja Agricola) of the Canarian Island Government in Arucas, Gran Canaria, Spain. In total, 45 female Canarian goats, weighing between 12-35 kg and no less than four months old, were purchased from a local dairy farmer and transported to the study site. The goats were placed in fly-proofed pens and allowed to acclimatise for two weeks, before being randomly selected and divided into test groups of four. Goats were experimentally infected intravenously from two highly parasitaemic donor goats, with 10^6^ and 10^5^ parasites per goat for *T. congolense* and *T. vivax*, respectively. AN7973 was administered intramuscularly accordingly, as either two injections of 10 mg/kg or as a single bolus dose of 10 mg/kg, on days 7 and 8 post-infection. Thereafter, the parasitaemia was monitored in the goats for up to 100 days post-treatment, after which any aparasitaemic and surviving goats were considered cured. Relapsed goats were removed immediately from the trial and humanely euthanized with an intravenous injection of sodium phenobarbital. The experimental protocols used for the goat studies were approved by the ethics committee for animal experimentation by the Veterinary Faculty of the University of Las Palmas in Gran Canaria, Spain on July 21, 2012 with the reference number 240/030/0121-36/2012. The studies were conducted under the strict guidelines set out by the FELASA for the correct implementation of animal care and experimentation.

The efficacy of AN 7973 against *T. congolense* and *T. vivax* in cattle was tested as described in [67]. The studies were conducted in fly-proof facilities and included negative (saline) controls; and the staff were blinded with regard to allocation of animals to treatment groups. Assessments were made for 100 days post treatment unless animals relapsed sooner. Tests were done in accordance with the principles of veterinary good clinical practice (http://www.vichsec.org/guidelines/pharmaceuticals/pharma-efficacy/good-clinical-practice.html). The ethical and animal welfare approval number was 00 l-2013/CE-CIRDES.

### Trypanosome culture and compound treatment

Bloodstream-form *T. brucei brucei* 427 Lister strain were cultured in HMI-9 plus 10% foetal calf serum or CMM [68] plus 20% foetal calf serum at 37°C, 5% CO_2_. PCF *T. b. brucei* were cultured in SDM79 plus 10% foetal calf serum at 28°C. Compounds were dissolved at 20 mM in DMSO and aliquoted to avoid excessive freeze-thaw cycles.

EC_50_s were measured in two different ways. To obtain the EC_50_s in S1 Fig, compounds were serially diluted over 24 doubling dilutions in 100 µL culture medium in 96 well opaque plates from a starting concentration of 100 µM. Bloodstream form trypanosomes were added at a final density of 2×10^4^/ml (100 µl) and incubated for 48 hours, while procyclic forms were added at a final density of 2×10^5^/ml and incubated for 72 hours. After incubation, 20 µL of 0.49 mM Resazurin sodium salt in PBS was added to each well and plates were incubated for a further 24 hours. Plates were read on a BMG FLUOstar OPTIMA microplate reader (BMG Labtech GmbH, Germany) with λ_excitation_ = 544 nm and λ_emission_ = 590 nm.

For the results in Table 1, bloodstream form trypanosomes were diluted to 4000/ml in the presence of compounds (diluted in water from DMSO) and incubated for 72h. 3-4h before the end of the incubation, Resazurin (Sigma) was added (final concentration 44 µM). Resazurin fluorescence was measured to assess the number of surviving viable cells [69, 70]. Each assay was performed with 3 technical and 3 biological replicates.

For the time to kill assay (Supplementary Figs S1 and S2) bloodstream-form *T. b. brucei* were cultured in 24 well plates in triplicate. Cultures were seeded at 5 × 10^5^/ml and compound was added at 5xEC_50_. Cells in each well were counted at 2, 4, 6, 8 and 24 hours using a haemocytometer.

For the all assays except the metabolomes shown in Fig 1 and S6 Table, compounds were used at 10x the 72-h EC_50_. To allow for variations between drug aliquots, EC_50_s were measured prior to every experimental series. The concentrations of compounds used in different experiments are listed in S1 Table and S6 Table.

### Cell cycle, protein and Northern blot analyses

For protein analysis, 2-3×10^6^ cells were collected per each sample, resuspended in Laemmli buffer heated and subjected to SDS-PAGE gel electrophoresis. All assays of macromolecular biosynthesis and RNA processing were done at densities of less than 2 × 10^6^/ml. Pulse-labeling was done as described in [71].

Total RNA was extracted from roughly 5×10^7^ cells using peqGold TriFast (peqLab) following the manufacturer’s instructions. The RNA was separated on formaldehyde gels and then blotted on Nytran membranes (GE Healthcare). Following crosslinking and methylene blue staining (SERVA), the northern blots were hybridized with the appropriate probes. For mRNA detection, the membranes were incubated with [α-^32^P]dCTP radioactively labelled DNA probes (Prime-IT RmT Random Primer Labelling Kit, Stratagene) overnight at 65°C. For spliced leader detection, a 39mer oligonucleotide complementary to the spliced leader was labelled with [γ-^32^P]ATP using T4 polynucleotide kinase (NEB) and incubated with the membrane overnight at 42°C. After washing the blot, it was exposed to autoradiography films and detection was performed with FLA-7000 (GE Healthcare). The images were processed with ImageJ.

### Primer extension assays

Compound treatments were all done at cell densities of about 0.9×10^6^ cells/ml. For each condition, 8-10×10^7^ cells were used. Primer extension was done approximately as described in [49]; primers were: ACCCCACCTTCCAGATTC for *SLRNA* (KW01 or CZ6364) and TGGTTATTTCTCATTTAAGAGG (CZ6491) for U3 snRNA. Both primers and the ladder were radioactively 5’-end-labelled with [γ-^32^P]ATP. For extension, 10 µg of RNA was incubated for 5’ at 65° with 2 µl of dNTPs (10 mM) and roughly 200 000 counts per minute (cpm) of the corresponding primer. Afterwards, RNasin (Promega), SuperScript^®^ III Reverse Transcriptase (Thermo Fischer), DTT and buffer were added according to the manufacturers instructions. The mixture was incubated 60’ at 50°C and then inactivated 15’ at 70°C. The samples were run in 35 cm long 6% polyacrylamide gels, dried, and analysed by phosphorimaging, The images were analysed using Fuji / ImageJ.

### In vitro transcription and splicing

In vitro transcription in permeabilised cells was done following the published procedure [50, 60] with minor modifications [72]. Briefly, cells (2.5 × 10^8^/reaction) were permeabilised with lysolecithin for 1 min on ice, washed, then resuspended in 60µl transcription buffer. 1 µl of either AN7973 or DMSO alone were added, and the reaction pre-incubated at 28°C for 2 min. After addition of 100µl transcription cocktail containing 1 µl of either AN7973 or DMSO, the reaction was allowed to proceed for 10 min at 28°C. The permeabilised cells were pelleted (45sec) and resuspended in 1 ml of TriFast. After the first phase separation, the aqueous fraction was re-extracted with phenol to remove residual protein; this was necessary to obtain good separation during gel electrophoresis. The final RNA volume was 20 µl. 10µl of each reaction were separated on a 7% polyadcrylamide/urea gel and the products were detected by phosphorimaging with low molecular weight DNA markers (New England Biolabs).

### Genomic sequence analysis

For genomic DNA sequencing, libraries were prepared at the Cell Networks Deep Sequencing Core Facility (University of Heidelberg) and subjected to paired-end MiSeq (Illumina) at EMBL. The quality of sequencing was evaluated with FastQC and the reads were trimmed using Trimmomatic. The output was aligned to the *T. b. brucei* TREU927 genome (version 9.0) using bowtie2. Results are available in ArrayExpress under accession number E-MTAB-6307. The Picard option AddOrReplaceReadGroups was used to create a valid .bam file to then be piped into GATK for obtaining, as output, .vcf files containing SNP and indel information. SnpSift filtered the features of interest, excluding for example synonymous mutations and intergenic regions. Identified variations from all cell lines were pooled to look for mutations found in all strains compared to the wild type, and in addition, reads from each strain was processed separately to find all the mutated genes. The lists were then compared. The genes taken into consideration were identified with annotation and categories from the Clayton lab in-house annotation list. Based on these annotation, the genes were filtered in Excel, where highly repetitive genes were excluded from the analysis. At first, variant surface glycoproteins (VSG), expression site-associated genes (ESAG), receptor-type adenylate cyclase GRESAG, UDP-Gal or UDP-GlcNAc-dependent glycosyltransferases and pseudogenes were removed. In the case of non-homozygous mutations, the genes were further selected filtering out other repetitive genes such as leucine-rich repeat proteins (LRRP), various invariant surface glycoproteins (ISG), adenosine transporters (TbNT) and retrotransposon hot spot (RHS) proteins. The list of gene IDs was compared with their translation level based on ribosome profiles [73, 74] and genes with values below 10 (non-translated) were excluded from the analysis.

### Metabolomics

For all assays at 5x EC_50_, bloodstream-form *T. b. brucei* were inoculated into medium at 1 × 10^6^/ml (for a six or eight hour incubation) or 2 × 10^5^/ml (for a 24 hour incubation) and compounds were added. Cells were incubated with compounds under normal growth conditions. At the desired time point, 1 × 10^8^ cells were taken and cooled rapidly in a dry-ice ethanol bath to 4°C. Samples were centrifuged at 1250 g twice to remove all medium before 200 µL chloroform:methanol:water (1:2:1) were added. Extracts were shaken for one hour at max speed before cell debris was removed by centrifugation at 16,000 g. Metabolite extracts were stored at −80°C under argon gas.

*Mass spectrometry –* Metabolite samples were defrosted and run on a pHILIC column coupled to an Orbitrap mass spectrometer as previously described [75]. In batch 1 an Orbitrap Exactive (Thermo Scientific) was used with settings including mass range: 70-1400, a lock mass of 74.0964, capillary: 40V, Tube lens: 70V, Skimmer: 20V, Gate lens: 6.75V and C-trap RF: 700V. In batch 2 an Orbitrap QExactive (Thermo Scientific) was used with settings including mass range: 70-1050, lock masses of 74.0964 and 88.0757, S-lens: 25V, Skimmer: 15V, Gate lens: 5.88V and C-trap RF: 700V. For fragmentation analysis an MS2 isolation window of 4 m/z, an intensity threshold of 3.3e5 and dynamic exclusion of 10 seconds were used.

Metabolites were putatively annotated using IDEOM software [76] before verification of annotations using mass, retention time, isotope distribution and fragmentation pattern. Xcalibur (Thermo) was used to explore the raw data, MzCloud (mzcloud.org) was used to match fragments to database spectra. Metabolite analysis was done using four biological replicates per condition and cell line. Relative metabolite levels were based on raw peak height relative to the average raw peak height of untreated cells. Lists of metabolites were mapped on metabolic pathways using Pathos (http://motif.gla.ac.uk/Pathos/). Intersections between samples were found using bioVenn (http://www.cmbi.ru.nl/cdd/biovenn/). (http://www.cmbi.ru.nl/cdd/biovenn/). Metabolite identities are consistent with standards from the Metabolomics Standards Initiative and evidence for each identity is shown in Table S6.

### YFP-DHH1 and expression of myc-tagged proteins

For over-expression of C-terminally myc-tagged proteins, the open reading frames encoding proteins of interest were amplified by PCR from genomic DNA, and cloned into pRPa-6xmyc [77]. After transfection, cells were selected and expression of myc-tagged protein was induced overnight with 100 ng/ml tetracycline.

The plasmid for creation of cells expressing YFP-DHH1 [78] was a kind gift from Susanne Krämer (University of Würzburg). It was transfected into bloodstream-form trypanosomes and two stable cell lines expressing the protein were selected. Trypanosomes (maximum density less than 1 million/ml in 10 ml) were treated for 30 min with 2 µg/ml Sinefungin, or for 1-2h with compounds at 10x EC_50_. After collection (5 min 1000g) and washing in PBS cells were resuspended in 20 µl PBS. 500 µl of 4% paraformaldehyde solution in PBS was added, cells were incubated without shaking for 18 min, washed 3x with PBS, then distributed onto poly-lysine coated chamber glass slides (all cells divided in two chambers) and left at 4°C over night. The PBS was then removed and cells were permeabilised using 0.2% Triton X-100 (w/v) in PBS, with shaking at room temperature for 20 min. After a further 3 washes, slides were incubate at room temperature with 200 ng/ml DAPI in PBS (15’ shaking), washed twice more, air-dried, embedded and covered for microscopy.

Slides were viewed and the images captured with Olympus IX81 microscope. The 100x oil objective was used. Digital imaging was done with ORCA-R2 digital CCD camera C10600 (Hamamatsu) and using the xcellence rt software. Bright field images were taken using differential interference contrast (DIC), Exposure time 30 ms, Lamp 4.0. Fluorescent images were made using DAPI and YFP filters. They were taken as Z-stacks, 30-40 images in a 8 µm thick layer, Exposure time 40 ms, Light intensity 100%, and afterwards they were deconvoluted (Numerical aperture 1.45, Wiener filter, Sub-Volume Overlap 20, Spherical Aberration Detection Accurate).

The fluorescence images were processed using ImageJ. For YFP and DAPI channels, layers containing signals ere selected, then the projection of maximum intensity of the deconvoluted stack was used. The images were saved in an 8-bit range then overlayed with DAPI in cyan and YFP as magenta. The colour balance was then adjusted with variable maxima for DAPI/cyan, but a set maximum of 80 for YFP/magenta.

After initial assessment after AN7973 and Sinefungin treatment, three independent experimental series, each including a negative control, were processed, and the images were read blinded. Cells were classified as having nuclear periphery granules if there were at least four strong granules on top of, or within one granule diameter of, the nucleus. Cells were classified as “strong” if nearly all granules were around the nucleus, and “possible” if there were several granules in other positions as well. For granule counts, only structures with at least four adjacent pixels at maximum intensity were considered.

## Acknowledgements

We thank Susanne Kramer (University of Würzburg, Germany) for the GFP-DHH1 plasmid and both Susanne and Shula Michaeli (Bar-Ilan University, Ramat-Gan, Israel) for useful suggestions and discussions. We thank Pieter Steketee and Julie Kovárová (University of Glasgow) for invaluable assistance with AN7973 metabolomics, and Ute Leibfried (ZMBH) for technical assistance. We also thank David Hepworth (Pfizer) for his assistance in manuscript clearance and checking compound structures.

## Supporting information

<u>S1 Table</u>

EC50s and compound concentrations used in different experimental series.

<u>**S2 Table**</u>

Copy number changes, single nucleotide polymorphisms (snps) and insertions and deletions (indels) in cells with weak resistance to AN7973

<u>**S3 Table**</u>

Details of snps and indels in cells with weak resistance to AN7973

<u>**S4 Table**</u>

Details of copy number variations in cells with weak resistance to AN7973

<u>**S5 Table**</u>

Metabolite changes after treatment of wild-type and mildly resistant cells with AN7973 for 5h.

<u>**S6 Table**</u>

Levels of chosen metabolites after treatment with different compounds. A detailed legend is on the first sheet.

<u>**S7 Table**</u>

EC50s of various benzoxaboroles in cells with and without additional CPSF3 expression.

<u>**S1 Fig**</u>

A. Growth curves of bloodstream-form *T. brucei* in the presence and absence of benzoxaboroles.

B. Structures of investigated benzoxaboroles, classified according to the time taken to affect growth (see (S2 Fig).

<u>**S2 Fig**</u>

Time-to-kill assays for benzoxaboroles - like S1 Fig but with more compounds and without the untreated control.

<u>**S3 Fig**</u>

Relationships between different measurements

(A) Relationship between the average increase in SAM and MTA and the percent Y structure in experiment 1.

(B) Metabolome of cells incubated for 6 h with either Actinomycin D (x-axis) or AN5568 (y-axis). More extensive results for AN5568 will been submitted separately.

(C) Relationship between the total number of perinuclear granules and the percent Y structure in experiment 1.

(D) The fold EC50 increase in cells expressing CPSF3-myc plotted against the percent Y structure in experiment 1.

<u>**S4 Fig**</u>

Effects of benzoxaboroles on Y structure levels.

A. Time course of effects of AN7973, AN3967, AN5827 and AN11736 in bloodstream forms (Experiment 2 in Fig 7). Sinefungin treatment was for 30 min.

B, C: effect of AN7973 in procyclic forms.

<u>**S5 Fig**</u>

Results for three separate experiments in which the location of YFP-DHH1 was analysed. Details are as in Fig 9B, but in addition, the average numbers of large granules (at least 4 contiguous pixels at maximum intensity) anywhere in the cell are displayed as black spots. The dotted line corresponds to the negative control.

<u>**S6 Fig**</u>

YFP-DHH1 localization after treatment with different benzoxaboroles. These are not “typical” images; instead, fields in which nuclear periphery granules were present have been chosen. Some (but not all) examples of peri-nuclear granule patterns are indicated by arrows. Compounds used are shown on each image. For AN7973 examples from the three different experiments are shown. The key is on the bottom left.

<u>**S7 Fig**</u>

YFP-DHH1 localization after treatment with different benzoxaboroles. These are not “typical” images; instead, fields in which nuclear periphery granules were present have been chosen. Compounds used are indicated and the key is as in S6 Fig. The control image in this case had brighter GFP fluorescence, for unknown reasons.

<u>**S8 Fig**</u>

Sequences of CPSF3 from *T. brucei* (TREU927), *Plasmodium falciparum* and *Homo sapiens*

Residues that were found to be mutated in AN3661-resistant *Toxopasma gondii* or *P. falciparum* are indicated with red arrows and the new residues are indicated in purple or cyan. The numbering is for *P. falciparum*. Black-highlighted residues are identical in at least two sequences and grey ones indicate conservative replacements.

<u>**S9 Fig**</u>

Details of EC_50_ results in cells with and without CPSF3-myc expression.

<u>**S10 Fig**</u>

Sequences of CPSF3 from *T. brucei* (TREU927), *T. vivax* and *T. congolense.* Details are as in S8 Fig.

